# Bistability of prefrontal states gates access to consciousness

**DOI:** 10.1101/2020.01.29.924928

**Authors:** Abhilash Dwarakanath, Vishal Kapoor, Joachim Werner, Shervin Safavi, Leonid A. Fedorov, Nikos K. Logothetis, Theofanis I. Panagiotaropoulos

**Author notes:** Equal contribution (Abhilash Dwarakanath, Vishal Kapoor). Co-senior authors (Nikos K. Logothetis, Theofanis I. Panagiotaropoulos).

## Abstract

Access of sensory information to consciousness is thought to be mediated through ignition of neural activity in the prefrontal cortex (PFC). Ignition occurs once activity elicited by sensory input crosses a threshold, which has been shown to depend on brain state fluctuations. However, the neural correlates of fluctuations and their interaction with the neural representations of conscious contents within the PFC remain largely unknown. To understand the role of prefrontal state fluctuations in conscious access, we combined multielectrode intracortical recordings with a no-report binocular rivalry (BR) paradigm that induces spontaneously-driven changes in conscious perception. During BR, antagonistic coupling of two prefrontal states, characterised by dominance of low frequency (1-9Hz) or beta (20-40Hz) local field potentials (LFP), reflect competition between two states of visual consciousness; perceptual update and stability, respectively. Low frequency perisynaptic bursts precede spontaneous transitions in conscious perception, signalling upcoming perceptual update of conscious content. We therefore show that it is a global cortical state that seems to drive internal switches, rather than the spiking activity of selective neuronal ensembles, which subsequently, only report the active percept. Beta band bursts were found to be correlated with periods of stable conscious perception, and selectively synchronised the neural ensemble coding for the consciously perceived stimulus. Similar ongoing fluctuations in the LFPs, with dynamics resembling the distribution of perceptual dominance periods during BR, dominated the prefrontal cortex during resting-state, thus pointing to their default, endogenous nature. Our results suggest that the two modes of conscious perception: perceptual update, and stability, can be associated with distinct prefrontal cortical states.

## Main Text

When the visual system is confronted with ambiguous sensory information, conscious perception spontaneously fluctuates between different possible perceptual interpretations (Leopold and Logothetis, 1999). In an unpredictable manner, one of the competing representations temporarily gains access to consciousness while the others become perceptually suppressed, therefore dissociating sensory input from subjective conscious perception. This perceptual multistability is a gateway in understanding the mechanism that allows the emergence of visual consciousness, due to the spontaneous switching of conscious perception between different co-registered representations. Although multistable perception has been intensively used to study the distribution of conscious content representations across the brain (Panagiotaropoulos et al, 2014), the exact neural mechanisms underlying the internally driven passage of sensory input from non-conscious processing to conscious access and vice versa remain largely unknown (Sterzer et al., 2009).

Binocular rivalry (BR) is a paradigm of multistable perception that is commonly used to identify this mechanism. During BR, the content of consciousness spontaneously alternates between two disparate stimuli that are continuously presented to each eye (Blake, 2001; Blake and Logothetis, 2002; Clifford, 2009). Two major mechanisms associated with different cortical areas have been hypothesised to drive these spontaneous transitions in conscious content: competition between monocular neurons in the primary visual cortex (V1), and activation of a widespread cortical network driven from a neural ignition event in the prefrontal cortex (PFC) (Blake, 1989; Dehaene et al., 2011; Mashour et al., 2020; Metzger et al., 2017). The originally-proposed mechanism that involves competition between monocular V1 neurons may not be sufficient to explain rivalrous switches in conscious perception (Leopold and Logothetis, 1996). This is because BR involves competition between higher-order perceptual representations that are not bound to eye-specific input (Logothetis et al., 1996), monocular neurons in V1 respond to the physical, but not the consciously perceived stimulus (Leopold and Logothetis, 1996) and switches in the activity of ocular dominance columns during BR can also be observed in V1 during anaesthesia (Xu et al., 2016). The second candidate mechanism involves higher cortical areas, in particular the PFC, (Leopold and Logothetis, 1999; Lumer et al., 1998; van Vugt et al., 2018) and suggests that fluctuations in cortical states may be critical for inducing ignition of neural activity, and therefore conscious access (Dehaene and Changeux, 2005; Doesburg et al., 2009; Levinson et al., 2021; Weilnhammer et al., 2021). Neural ignition associated with conscious perception manifests itself as a sudden and sustained non-linear increase in gross-scale activity measured by non-invasive, whole-brain recording techniques using electroencephalography (EEG), electrocorticography (ECoG) and magnetoencephalography (MEG), around 200-300ms after stimulus onset (Dehaene and Changeux, 2011; Dehaene et al., 2001; Del Cul et al., 2007; Fisch et al., 2009; Joglekar et al., 2018; Noy et al., 2015).

This concept of ignition, thought to be critical to conscious experience, falls under the purview of the Global Neuronal Workspace theory (GNW), which posits that access to consciousness is gated by non-linear amplification of neuronal activity in the PFC, and is broadcast to widely-distributed cortical networks. Perhaps the most persuasive evidence at the neuronal level for this theory comes from two recent studies, both leveraging the benefit of direct intracortical, electrophysiological recordings, and showing sustained and increased activity in the frontal regions, somatosensory and ventral premotor cortices at the single neuronal level when a percept enters consciousness (Noel et al., 2019; van Vugt et al., 2018). Particularly, the study by van Vugt et. al. reveals that the propagation of stimulus-driven activity from lower visual areas to the PFC, is influenced by pre-stimulus states, determining the conscious accessibility of sensory input. This initiates neural ignition, which is considered to be the neural correlate of threshold-crossing (Kang et al., 2017) for the detection of a stimulus in signal detection theory (SDT). Intuitively, breaching can also be understood in terms of a gating mechanism, an operation that could control the threshold to conscious perception in the PFC through the bistability of intrinsically-generated cortical states (O’Reilly, 2006). However, to date, there is scant evidence as to what neural processes are related to this frontal ignition mechanism, or the substrates of gating, that enable this access to consciousness.

In the present study, we attempted to unravel the mechanisms underlying the emergence of conscious perception in the macaque lateral PFC using a no-report BR paradigm. This allowed us to detect internally-driven transitions in the conscious perception of stimuli that moved in opposing directions. We combined this task with multielectrode recordings of local field potentials (LFPs), and simultaneously-sampled, direction-of-motion selective, spiking activity of competing neuronal ensembles. By using the optokinetic nystagmus (OKN) reflex as an objective criterion of perceptual state transitions, we removed any effects of voluntary motor reports on neural activity, thus identifying signals directly related to spontaneous transitions in the content of consciousness. Our results suggest that ongoing alternations in the cortical state between low-frequency (1-9Hz) and beta (20-40Hz) oscillations directly relate to update and stability of neural representations of conscious contents, respectively. Finally, we extend the concept of neural ignition to demonstrate that at the mesoscopic scale, a linear increase in the power of low-frequency activity coupled with a non-linear and sudden increment in neuronal nodes recruited in the PFC is critical in managing conscious access.

## Results

We used a no-report paradigm of binocular motion rivalry coupled with multielectrode extracellular recordings of LFPs and direction-of-motion-selective neuronal ensembles in the inferior convexity of the macaque PFC (Fig. 1A). Two trial conditions were employed: a) physical alternation (PA) of monocularly alternating gratings with opposing directions of motion and b) binocular rivalry (BR), where the initial direction of motion stimulus was not removed, but was followed by a grating moving in the opposite direction, presented to the contralateral eye (Fig. 1B, upper panel). This manipulation results in an externally-induced period of perceptual suppression of variable duration for the first stimulus (binocular flash suppression - BFS), which is then followed by spontaneous perceptual transitions, since the two competing representations start to rival for access to consciousness. In order to exclude the effect of voluntary perceptual reports on neural activity, the macaques were not trained to report their percept. Instead, the polarity of their motion-induced OKN elicited during passive observation of the stimuli (in both conditions, i.e. BR and PA), which was previously shown to provide an accurate perceptual state read-out in both humans and macaques (Fox et al., 1975; Logothetis and Schall, 1990), was used to infer perceptual dominance periods (Fig. 1B, lower panel). These dominance durations followed a gamma distribution, a hallmark of multistable perception, with a median dominance duration of 1.54±1.28s (median±SD) for spontaneous transitions in BR and 2.25±2.21s for transitions involving exogenous perceptual suppression in BFS (Fig. 1C).

**Figure 1.**
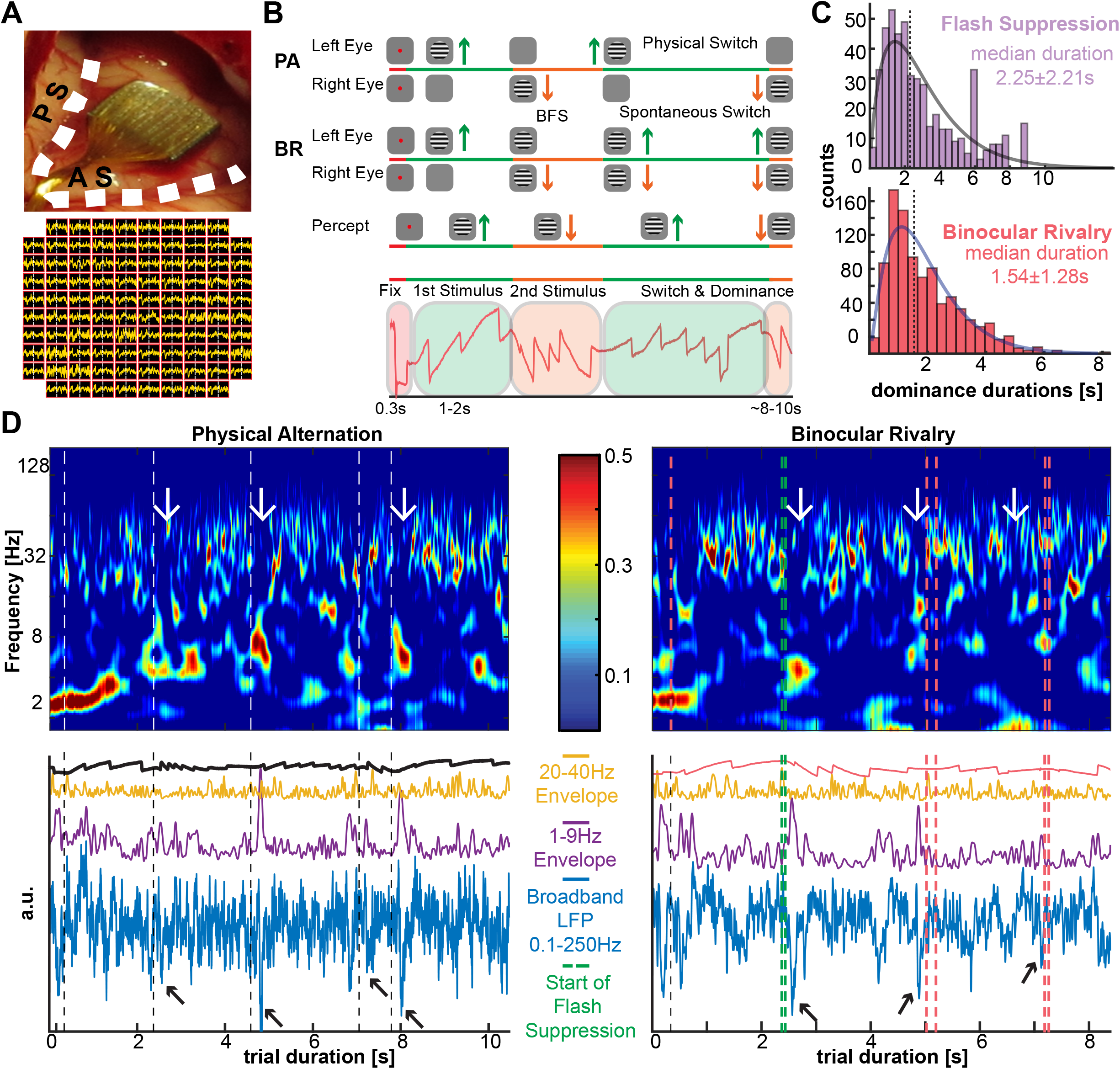
Experiment and typical perisynaptic signals. A. Multielectrode array in the inferior convexity of the PFC (top). AS: arcuate sulcus; PS: principal sulcus. Below, an example spatial map of 0.1-250 Hz LFP signals around a spontaneous (white line) perceptual transition. B. Task structure: For both PA and BR trials, following binocular fusion and fixation of a red dot (0.2°) for 300ms, a grating (or 200 random dots at 100% coherence in one session) moving upward (90°, green) or downward (270°, orange) was initially presented in one eye (8°, 12-13°/sec, 100% contrast). After 1000-2000 ms the first stimulus was removed in PA trials and followed by the presentation of the opposing motion direction and subsequently multiple monocular switches up to 8-10 seconds. In BR trials, after the presentation of the opposing motion direction (resulting in BFS) the two stimuli were let to compete for conscious access resulting in spontaneous perceptual switches. Different OKN patterns (red trace, highlighted in green and orange) elicited from two directions of motion allowed decoding of the conscious percept. C. Histogram and gamma distribution function fit of perceptual dominance times during BFS (2.25±2.21s, median±SD) and spontaneous perceptual transitions during BR (1.54±1.28s, median±SD). D. Channel-averaged and normalised (z-scored) time-frequency spectrograms for a single trial/observation period of PA (left) and BR (right) are shown at the top. White lines in PA reflect the manually marked change in the OKN polarity after the onset of the exogenous, monocular stimulus changes. Green lines in BR represent the start of the flash suppression phase (at 2.3sec) whereas red lines represent the subsequent spontaneous perceptual transitions. 1-9 Hz bursts suppressing 20-40 Hz activity occurred following a switch in PA but before a transition in BR (white arrows). Bottom panels: Broadband LFP, instantaneous 1-9 Hz and 20-40 Hz signal amplitude and OKN traces for the same observation period. Black arrows point to negative deflections in the 0.1-250 Hz LFP trace (for display, all traces were normalised and plotted with an arbitrary shift to clearly delineate the different oscillatory regimes).

### Prefrontal state fluctuations precede conscious perception

To investigate the role of internal oscillatory states in conscious refresh and update, we first analysed the perisynaptic activity dynamics, reflected in the LFPs, to identify mesoscopic signal fluctuations around spontaneous perceptual transitions, (Fig. 1D, Fig. S1). Transient negative deflections of the channel-averaged, raw LFPs (0.1-250 Hz, blue traces in Fig. 1D), were observed to disrupt a default state of oscillatory bursts in the beta range (20-40 Hz) throughout the observation periods in both PA and BR trials. However, the strongest negative deflections appeared to occur just after the change in the OKN polarity, induced by external stimulus changes in PA (Fig. 1D, left), but just before spontaneous perceptual transitions in BR (Fig. 1D, right). Pooling all physical stimulus transitions in PA (n=1322) revealed that the power of these transient negative deflections was concentrated immediately after exogenous stimulus changes in a low-frequency (1-9 Hz) range that resulted in a temporally transient suppression of the ongoing beta activity (20-40 Hz) (Fig. 2A, left).

**Figure 2.**
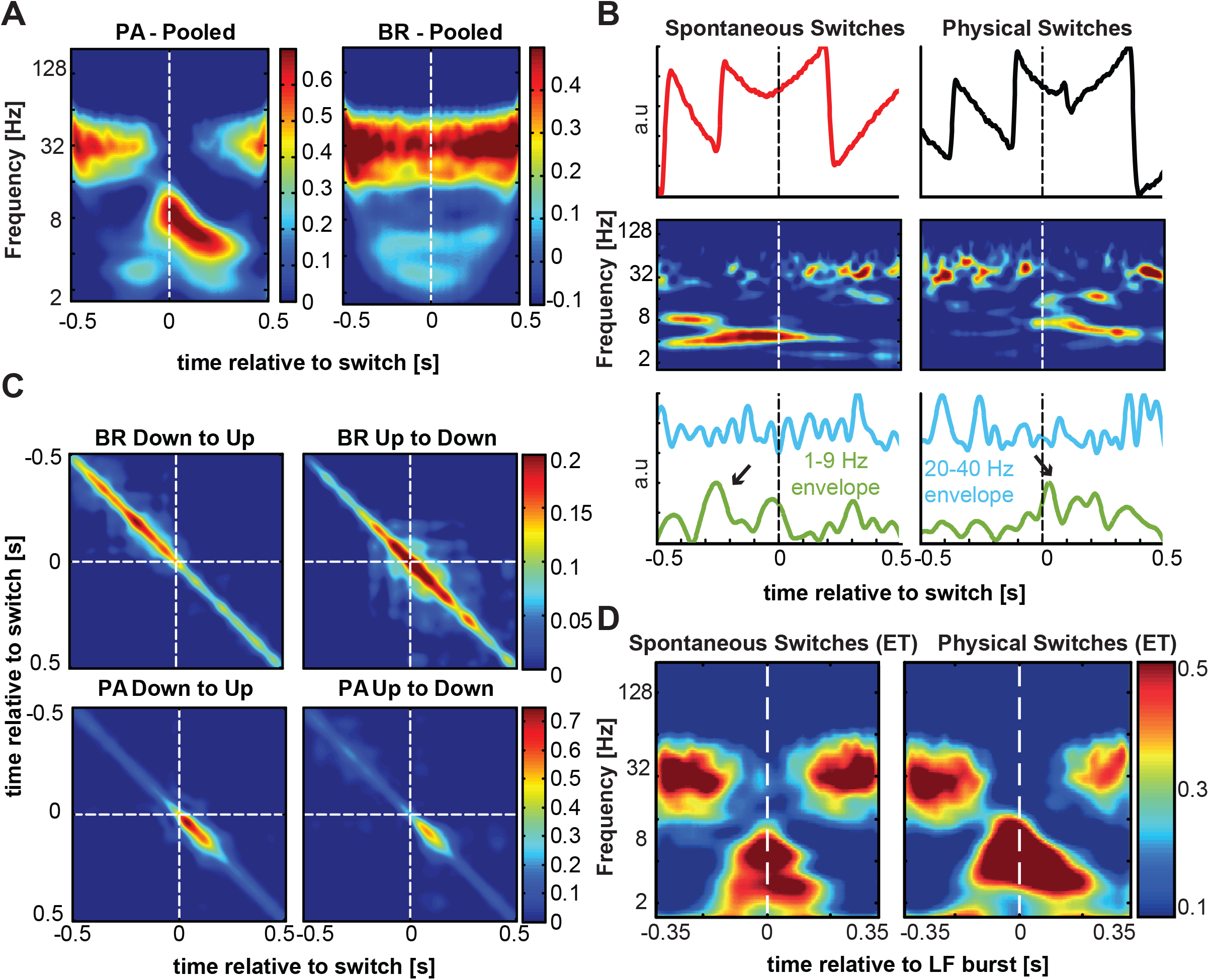
Time-frequency LFP analysis. A. Grand average time-frequency analysis of spontaneous transitions (right) and an equal number of randomly sampled physical (left) and transitions. Spectrograms are aligned (t=0) to manually-marked OKN changes in both PA and BR for periods of stable perceptual dominance before and after the switch. B. Upper panel: OKN traces around a single physical (black) and spontaneous (red) transition. Middle panel: channel-averaged normalised spectrograms aligned to the OKN slope change for the two conditions. Lower panel: Normalised instantaneous amplitudes of the two modulated frequency bands (i.e. 1-9 Hz, green and 20-40 Hz, cyan) identified from the spectrograms. Low-frequency bursts occur after the physical switch but before the spontaneous transition (black arrows). C. Differences in the onset of the low-frequency activity across physical and spontaneous transitions in direction of motion are reflected in the temporal auto-covariance of the low frequency envelopes across the array recorded simultaneously for every transition type (down to up, left and up to down, right). Most of the similarity is observed before a transition in BR (upper panel) but after a transition in PA (bottom panel). D. Grand-averaged LFP spectrograms aligned to low-frequency events, after a physical and before a spontaneous transition. The emergence of a low-frequency transient results in beta power suppression for around 300ms in both conditions.

Low-frequency-associated beta suppression was also observed for intrinsically-generated perceptual transitions in BR (n=573); however, it started well before (∼ 400ms) the spontaneous OKN change (Fig. 2B, right). In BR, the absence of a feedforward response, locked to an external change of the sensory input as in PA, resulted in a temporal jitter of the low-frequency transients across different transitions and neuronal sites (Fig. S2). In individual transitions, the low-frequency-associated beta suppression started clearly before the spontaneous OKN transition (Fig. 2B). Indeed, the temporal covariance of the channel-averaged low-frequency signal across transitions was concentrated well before the perceptual change in BR, whereas it was heavily concentrated in the post-transition period during PA (Fig. 2C). Spectrograms aligned to the low-frequency event peaks detected before and after the transition in BR and PA respectively show similarity in the coupling of low-frequency transients and beta-burst suppression between the two conditions (Fig. 2D). Therefore, these results show that suppression of 20-40 Hz activity during 1-9Hz transients follows exogenous stimulus changes in PA, but precedes spontaneous OKN - inferred perceptual transitions in BR.

Since both the low-frequency and beta activity were not sustained oscillations but appeared to occur in bursts, we quantified the burst-rate of low-frequency and beta activity, before and after the time of exogenous (PA) and endogenous (BR) perceptual transitions using a burst-rate metric (described in methods; only the burst-rates for low-frequency are reported). Low frequency burst rate (events/transition/channel) was significantly higher after the OKN change in PA (0.36±0.0046, n = 46495, post-transition, vs. 0.09±0.0014, pre-transition, n = 11330; p<10^−187^ mean ± SEM), but before the OKN change in BR (0.17 ± 0.002, n = 9670, pre-transition vs 0.14 ± 0.002, n = 7734, post-transition, p<10^−43^ mean ± SEM) (Fig. 3A). Furthermore, the low-frequency bursts were significantly more before a spontaneous perceptual transition than before a physical transition (0.17 ± 0.002, n = 9670, pre-transition BR vs. 0.09 ± 0.0014, n = 11330, pre-transition PA; p<10^−150^ mean ± SEM). To understand the average burst time, one must consider the fact that the post-switch period of the current transition, is the pre-switch period of the subsequent one. Therefore, we discarded events occurring after 250s, as this time period also encompasses the VEP time in PA transitions. Thus, low-frequency bursts occurred at around −114 ± 190ms median±SD. Importantly, low-frequency bursts, including the last-detected burst before a transition, occurred on average even before the end of the last dominance period preceding a transition in BR (end of dominance: −97.4 ± 140ms, low-frequency bursts collected up to the beginning of the next transition: −198 ± 133ms, median±SD, p<10^−67^, see Fig. 3B, and Fig. 2B for an example transition). As expected, low frequency bursts occurred predominantly and significantly after the OKN change in PA (64ms ± 147ms, median±SD). When the switches were aligned to the experimental TTL pulse, these bursting times were further shifted (190 ± 162ms, median±SD). This happens because the stimulus changes first, followed by the change in the OKN, which we consider as the exact switch-time (Fig. S3)

**Figure 3.**
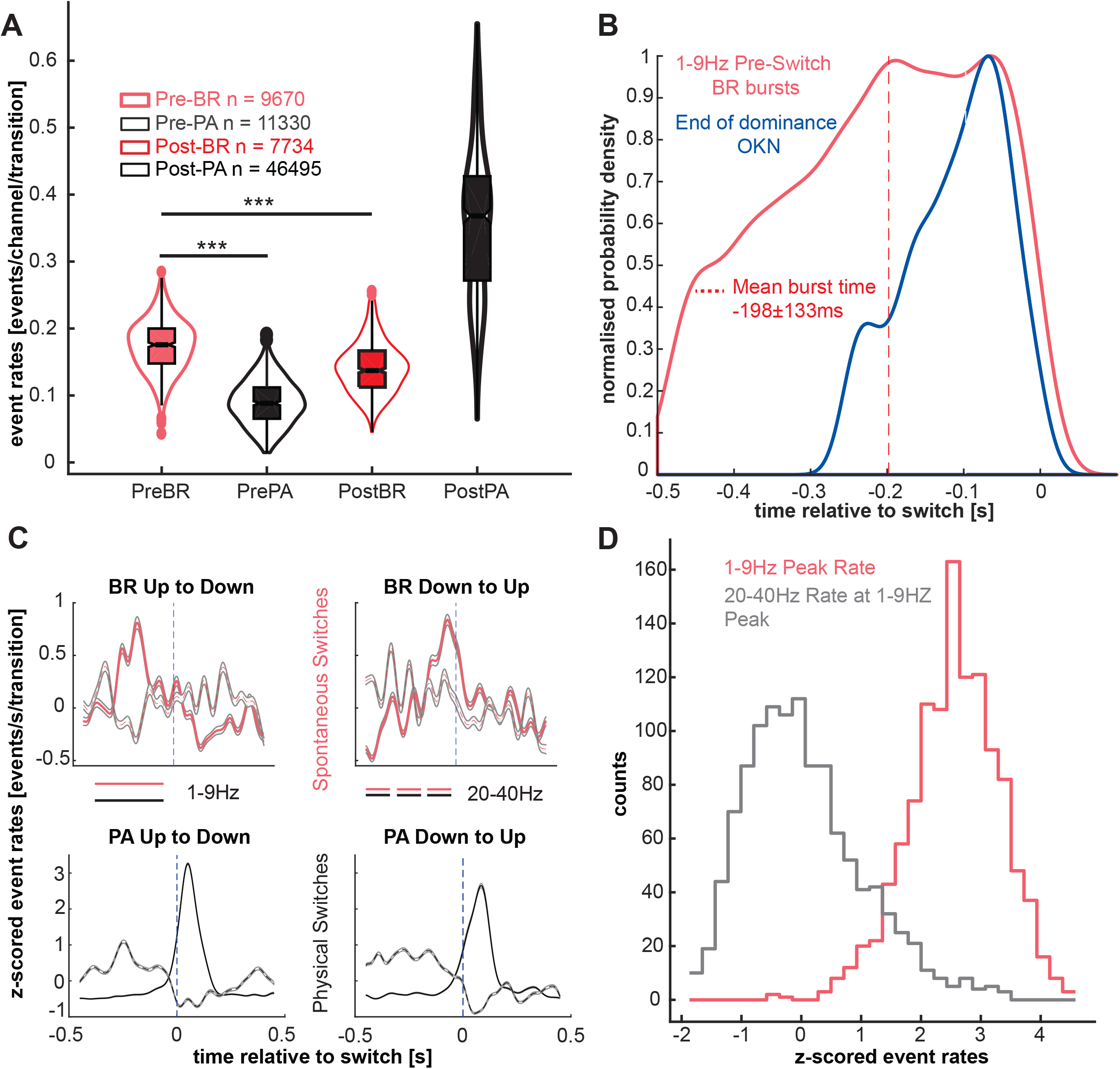
LFP burst-rate analysis. A. Burst-rate per transition per channel before and after transitions in PA (red) and BR (black). In order from left to right: Pre-BR (dark pink), Pre-PA (black), Post-BR (light pink), Post-PA (grey). The whiskers of the box-plots show the dispersion of the data. Notches depict median values. Matching colour dots represent outliers. More low frequency bursts occur before spontaneous, but after physical switches. Burst-rate before a physical switch is low, suggesting noise levels. This baseline burst-rate needs to be ramped up for a switch to occur. PA: (0.36 ± 0.0046, n = 46495, post-transition, vs. 0.09 ± 0.0014, pre-transition, n = 11330; p<10^−185^ mean ± SEM) BR: (0.17 ± 0.0016, n = 9670, pre-transition vs 0.14 ± 0.0016, n = 7734, post-transition, p<10^−43^ mean ± SEM). B. Distribution of low frequency burst-times and OKN times marking the end of the previous dominance period before a spontaneous transition. We fit the probability distribution using a kernel density estimate with a variable width. These functions were then normalised for direct comparison. Low-frequency bursts on average occurred at −198±133ms before a switch (red dashed line), significantly before the end of dominance which occurred at −97.4±140ms before a transition to the competing representation (p<10^−67^ median±SD). C. Normalised (z-score) burst-rate in time (events/s/transition) during BR (red lines) and PA trials (white lines) for low-frequency (solid lines) and beta activity (dashed lines). D. Distribution of beta-band rate (gray) at the peak of low-frequency rate (pink) before spontaneous perceptual transitions in BR. (r = −0.08, p=0.0071; pooled across both transition types)

The low-frequency activation before a spontaneous perceptual reversal in BR is better observed in the evolution of bursting activity in time (quasi-PSTH, i.e. the detected bursts are converted into binary event trains, smoothed and then averaged, i.e. events/s/transition). In BR, the peak-rate of 1-9 Hz bursts occurred at −160 ± 237ms and −28 ± 199ms (median±SD) before the spontaneous perceptual transitions for the two transition types respectively (Figure 3C, top row), while in PA they occurred at 52 ± 28ms (median±SD) and 82 ± 64.5ms (median±SD) following the marked OKN change (Fig. 3C, bottom row). These differences were further enhanced when the bursts towards the end of the post-switch window (i.e. bursts occurring 250ms after the transition, encompasses the average VEP in physical alternation trials, and considers that these events would be pre-switch events of the next switch) were discarded (Fig. S4). Confirming the time-frequency analysis pattern in Fig. 2A and suggesting a frequency-specific competitive process (i.e. cortical state fluctuations) in the PFC, the low-frequency and beta burst-rates were significantly anti-correlated in BR (r = −0.08, p = 0.0071; pooled across both transition types; Fig. 3D).

### Spatiotemporal build-up of prefrontal activity

Are the perisynaptic transients preceding a spontaneous change in the content of consciousness, random, large excursions from baseline activity, or do they reflect a gradual, spatiotemporal build-up process that is critical for inducing a spontaneous transition? Indeed, we noticed that in many instances before a spontaneous transition, the last transient low-frequency (1-9 Hz) burst was frequently preceded by similar but of lower amplitude bursts (Fig. 1D and Figure S1). When the instantaneous amplitude of the low-frequency activity triggered at every switch was averaged first across channels and then across transitions, we observed a gradual increase approaching spontaneous but not physical transitions (Fig. 4A). Fitting a line to the relationship between the transition-averaged low frequency burst amplitudes at every time point before a transition revealed a linearly increasing relationship between the two variables before a spontaneous (adjusted R^2^ = 0.34) but not before a physical switch (adjusted R^2^ = −0.003) (Fig. 4B). While this low-frequency burst amplitude exhibited a gradual linear increase, the number of activated prefrontal sites abruptly increased just before a spontaneous reversal, suggesting a non-linear increase in the spatial spread of prefrontal activation just before a spontaneous change in the content of consciousness (Fig. 4C and Figure S5). Taken together, these results indicate the occurrence of a mesoscopic, spatio-temporal spread of low-frequency prefrontal bursts before spontaneous perceptual reversals. Both linear and non-linear increases in the amplitude and spatial spread of activation, respectively indicate the operation of both modes in a prefrontal ignition process (Moutard et al., 2015) during BR.

**Figure 4.**
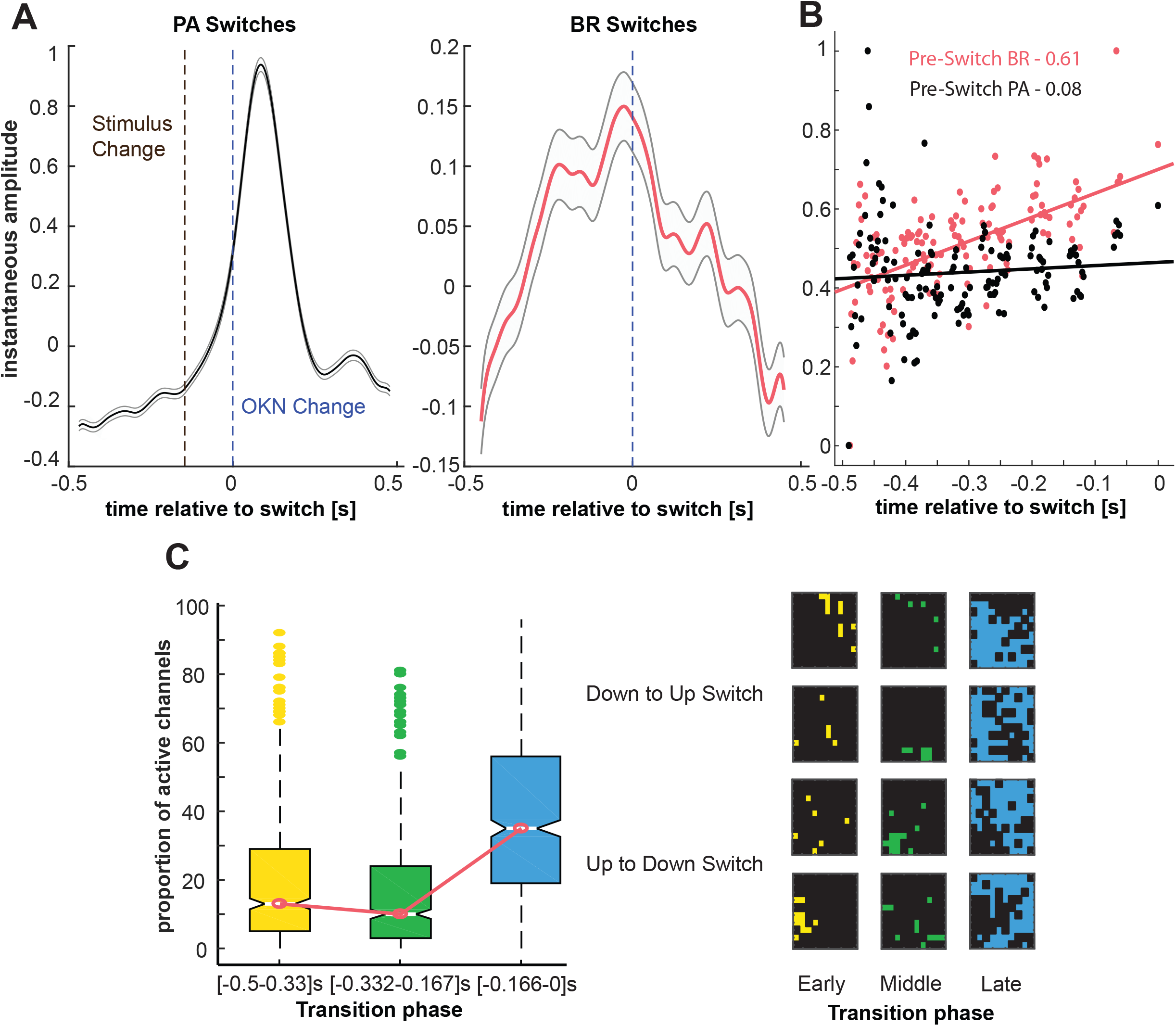
Spatiotemporal build-up preceding spontaneous transitions. A. Low-frequency instantaneous amplitude shows a slow climbing activity before a perceptual transition (right) but not before a physical transition (left). Curves reflect an average ± SEM across transitions of the channel-averaged activity for each collected transition. The black dashed line represents the onset of the stimulus detected by a TTL pulse. B. Average build-up of low-frequency activity in time is analysed by fitting a linear model to the pooled, and averaged amplitude at every time-point (red – BR, black – PA) across all measured neuronal sites. While before a spontaneous transition, low-frequency activity ramps up in time (slope = 0.61, R^2^ = 0.34), before a physical transition, it remains flat (slope = 0.08, R^2^ = −0.003). The instantaneous amplitudes here are normalised at the time of curve fitting. C. The spatio-temporal spread of low frequency activity is shown as the proportion of channels for a given transition that displayed low-frequency burst peaks in early ([−0.5 to −0.333s]), middle ([−0.332 to −0.167s]) and late temporal windows ([−0.166 to 0s]). A significantly larger proportion (p<0.01) of neuronal sites peak closer to the switch, showing a sharp, non-linear increase (left). Activation of sites on the array shown for four examples of different types of spontaneous transitions from 2 macaques (right). Dots indicate outliers.

We further hypothesised that if an increase in low-frequency bursting is critical for inducing spontaneous perceptual reversals, then low-frequency amplitude should be significantly weaker when perceptual transitions were not complete, but resulted in piecemeal (PM) periods in which perception did not unambiguously favour either of the two competing directions of motion (Fig. 5A). Subtracting the time-frequency decomposition of transitions to a PM percept from that of clean BR perceptual transitions revealed a preponderance of low-frequency activity before a switch, suggesting that the amplitude of low frequency bursts is indeed critical for completing a perceptual transition to another period of clear dominance

**Figure 5.**
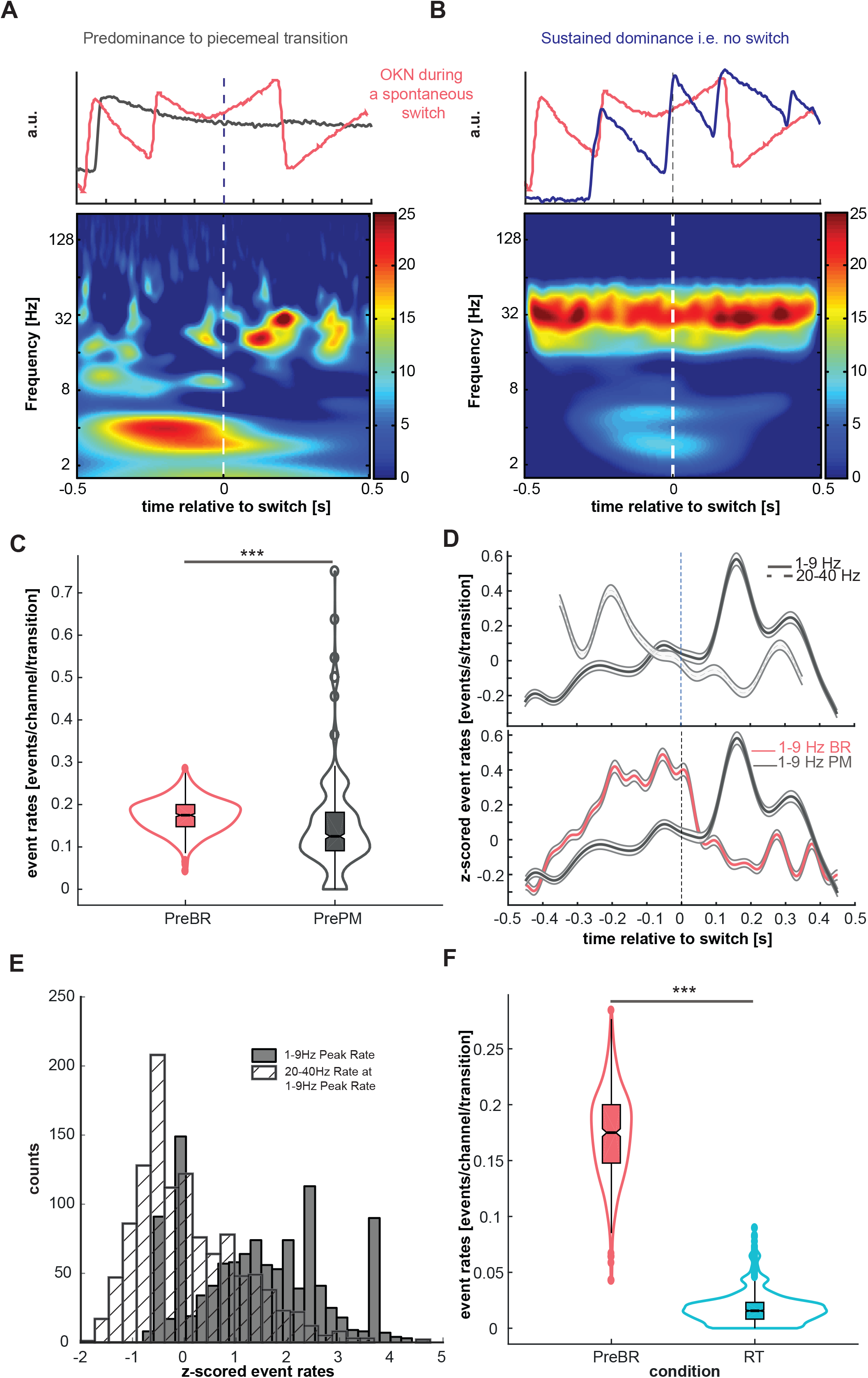
Low frequency amplitude and burst rate is critical for inducing clear switches. A. Top panel shows two typical OKN patterns elicited during a spontaneous transition (red) and during a transition to piecemeal (grey). The subtracted spectrogram (BR minus PM) below shows a large difference in low-frequency activity before a spontaneous switch suggesting very weak low frequency activation during a transition to piecemeal perception. B. Top panel shows two typical OKN patterns elicited during a spontaneous transition (red) and during sustained dominance where no switches occurred (blue). The subtracted spectrogram (BR minus no switches) recovers the time-frequency pattern observed during BR suggesting that the LFP activity during sustained perceptual dominance periods is at noise level. C. The low frequency burst rate per transition per channel before a spontaneous transition (Pre-BR, n= 9670, 0.17±0.0016) when compared to the period before transition to the piecemeal percept (Pre-PM, n= 2486, 0.147±0.004) is significantly higher (p < 10^−26^). D. Top row panel: z-scored event rates in time (events/transition/s) before and after the transition to piecemeal percepts, the solid line being the low-frequency component and the dashed line being the beta-band activity. While before the transition to piecemeal, the beta dominates, signalling the active percept, after the transition the low-frequency inhibits the beta thus signalling an upcoming percept. Bottom row panels: z-scored event rates in time (events/transition) for the low frequency activity for BR (red) and PM (grey). The low frequency activity burst rate peaks before transition to another clear dominance during BR (red) whereas it peaks after the transition to a piecemeal percept (grey). E. The distribution of the peak low-frequency rates vis a vis the rate of the beta activity at the timing of the low-frequency peak reveals no significant antagonism before a transition to a piecemeal (r = −0.0073, p=0.805) as compared to before clear spontaneous transitions, where a significant decoupling is observed (r = −0.08, p=0.0071). F. Low-frequency event rate per transition per channel for the two conditions, spontaneous switch and randomly triggered periods during sustained dominance. The burst rate before a spontaneous switch (pre-BR, 0.17±0.0016, n = 9670) is significantly higher than during sustained dominance (SD, 0.015±0.0005, n = 55026, 100 iterations, 550.26 bursts per iteration).

Next, we sought to clarify if there is a difference between low-frequency activity leading to a perceptual transition compared to low-frequency activity when perceptual transitions don’t occur, i.e. during periods of sustained dominance. To accomplish this, collect LFP activity around randomly-triggered time points during predominance. Subtracting the time-frequency decomposition of these randomly-triggered (RT) periods from that of BR, preserved the pattern observed in the latter (Fig. 5B), indicating that weak, low-frequency activity occurs as baseline noise, which only leads to a perceptual change when it is spatiotemporally ramped up in a structured manner (Figure 4B vs Figure 2A & Figure 3B). Indeed, we computed a mean rate of 0.015±0.0005 (n = 55026 after resampling 100 times, i.e. only 550.26 bursts per iteration) bursts per randomly triggered period during sustained dominance; an order of magnitude lower than the corresponding periods during BR (0.17±0.0016, n = 9670, events/transition/channel, Figure 5C). Furthermore, the proportion of sites that displayed low-frequency bursting activity across all BR periods was 100%, compared to only 51% during sustained dominance. These results further indicate that an increase in low-frequency burst-rate, build-up, and a larger spatial spread of low-frequency activation is necessary to drive spontaneous transitions. Additionally, the low-frequency burst-rate was higher before a clear spontaneous transition compared to the period before transition to a piecemeal percept (0.17±0.0016, n = 9670, pre-BR, vs. 0.14±0.004, pre-PM, n = 2486, events/transition/channel; p<10^−26^) (Figure 5D), with the low-frequency peak rate occurring after the transition to piecemeal (Figure 5E). Moreover, the burst rate was significantly higher after the transition to a piecemeal period (0.16±0.004, PM, n = 3531, vs 0.14±0.004 pre-PM, n = 2486, events/transition/channel; p<10^−5^), while the anti-correlation between low-frequency and beta was significant but weaker compared to clear spontaneous transitions (r = −0.009, p<10^−137^ vs r = −0.05, p<10^−295^, p < 10^−14^, Figure 3D and Figure 5F).

These results suggest that low-frequency activity should be significantly up-modulated from noise level, potentially crossing a threshold, to induce a perceptual transition.

### Conscious content-specific ensemble activity transitions and LFPs

We also investigated whether the LFP bursts precede the change in the encoding of the active conscious percept from the spiking activity of neuronal populations, which could shed light on the relationship between the global oscillatory states and ensemble activity. To understand this temporal relationship between perisynaptic state fluctuations and neuronal populations reflecting the content of consciousness, we compared the convergence times of the normalised discharge activity of simultaneously recorded ensembles selective for the rivalling gratings (i.e. the point at which the stimulus-correlated firing rate of the ensembles selective to the dominant and suppressed stimuli flips), (see Methods), Fig. 6A, Fig. S6), with the low-frequency burst and peak-rate distributions. We found that discharge activity converged significantly later compared to both the low-frequency peak-rates (−60 ± 222ms for LFP event/s/transition vs. 209 ± 295ms median±SD, convergence of spiking; p<10^−47^) and burst-times (−114 ± 190ms median burst time (truncated at 250ms post-switch) vs. 209 ± 295ms, median±SD convergence of spiking, p<10^−94^) in spontaneous perceptual transitions. In the majority of spontaneous transitions, spiking activity crossovers occurred after the median truncated low-frequency peak-rates and burst-times (in 86.2% of trials compared to peak-rates and in 89% of trials compared to burst times; Fig. 6B, Fig. S3). Therefore, low-frequency transients could reflect a pre-conscious process preceding an intrinsically generated perceptual transition.

**Figure 6.**
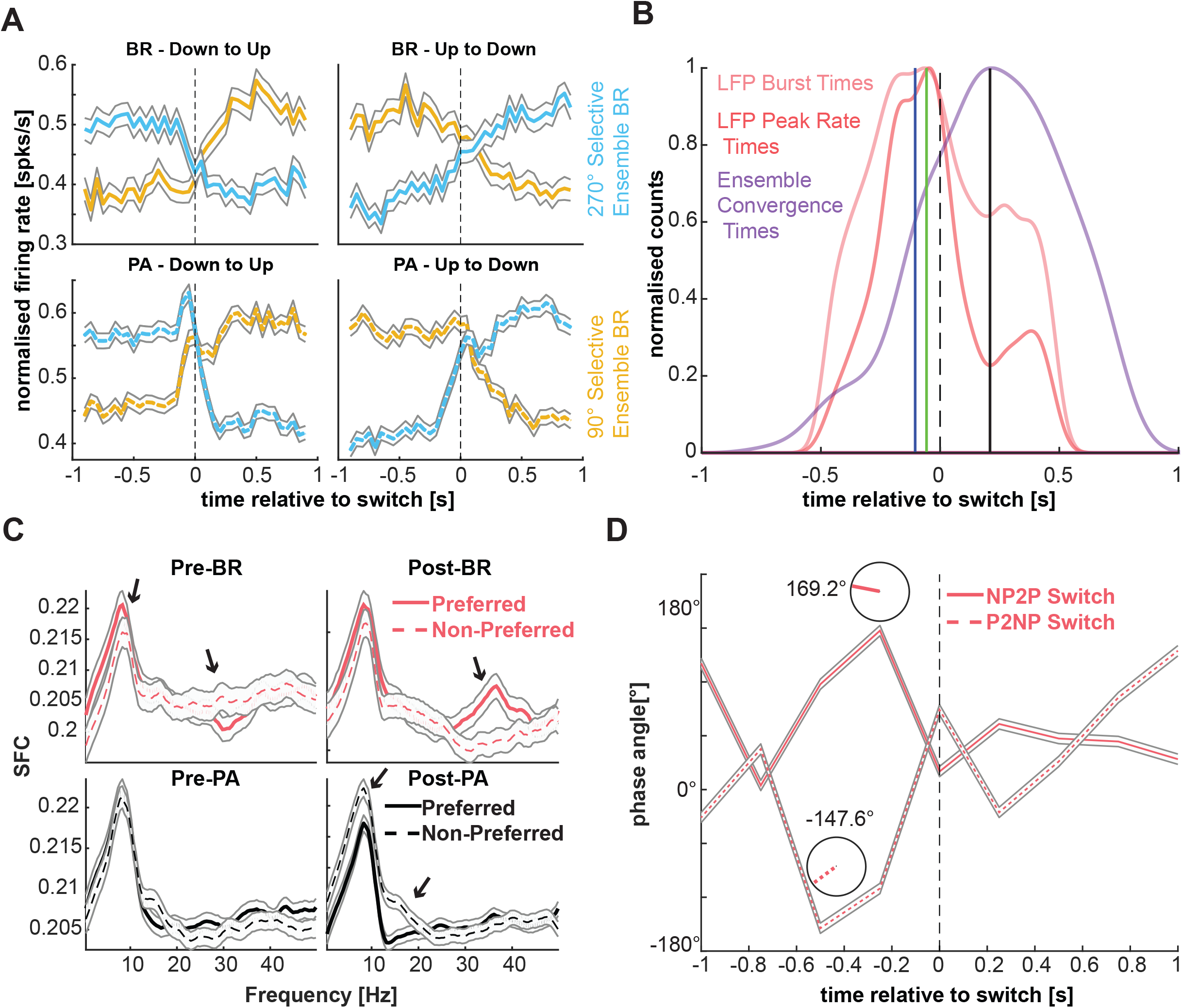
Relationship between the spiking activity of selective ensembles and the LFP. A. Top row: Average (across all collected transitions) population PSTHs of simultaneously recorded feature specific neuronal ensembles during BR. For each transition, we summed spiking activity across all selective single neurons belonging to each competing ensemble (upward and downward selective, blue and yellow, respectively), and normalised them. Bottom row: same for PA. B. Distribution of the (truncated) low-frequency burst times (pink), event-rate peak times (red) and the time of convergence of the PSTHs of each competing ensemble for each transition estimated by LOWESS smoothing and interpolation. Low-frequency activation preceded a change in the content of consciousness from neuronal populations (−60 ± 222ms for LFP event/s/transition (green line) and −114 ± 190ms (blue line) vs. 209 ± 295ms median±SD (black line), convergence of spiking; p<0.05 for both comparisons). D. After a spontaneous transition, spiking of the dominant population was significantly more coherent with the beta LFP band, as compared to the suppressed population (black arrows point to frequency bins with statistically significant differences between dominant and suppressed p<0.05). SFC during pre-switch BR periods when low-frequency transient bursts are more prevalent did not exhibit similar differences in the beta band. These effects in the beta band were absent in physical transitions where SFC for a dominant preferred stimulus was significantly reduced in a lower frequency range. E. Mean angles of spike-LFP phase locking during a non-preferred to preferred transition (“NP2P”, dashed line) and preferred to non-preferred transition (“P2NP”, solid line). The populations selective to the suppressed stimulus before a switch progressively become locked to the depolarising phase of the low-frequency LFP (169.2°), thereby causing them to increase their firing rate when their preferred stimulus becomes dominant after the perceptual transition.

Finally, to understand how prefrontal state fluctuations may be related to spiking-network reorganisation and therefore perceptual update, we computed the spike-field coherence, in a 500ms window preceding and succeeding a perceptual switch, (SFC) (Chandrasekaran et al., 2009; van der Meer and Redish, 2009) of the simultaneously recorded, feature-selective ensemble activity and the global broadband LFP across all transitions. After a spontaneous perceptual transition in BR, when the negative LFP deflections and therefore the low frequency (1-9 Hz) transients were less prevalent, the perceptually dominant ensemble was more coherent in the beta range (∼25-37 Hz) compared to the suppressed ensemble (p < 0.03; Fig. 6C). However, there were no differences between the suppressed and dominant populations in the period approaching a spontaneous transition when low frequency transients accompany a suppression of beta bursts. These results suggest that the prefrontal ensemble signalling the current content of consciousness is synchronised in the beta-band of the LFP. Low-frequency transients dissolve the beta-coherent ensemble, potentially increasing the likelihood for spiking in the suppressed population and therefore the likelihood for perceptual reorganisation. Further evidence for this hypothesis is seen in the mean phase-angles of spike-LFP coupling in the low-frequency band (Figure 6D). Before a switch, sites that prefer the suppressed stimulus (non-preferred to preferred switch; NP2P) are locked to the depolarising phase of the LFP (169.2°), while sites that prefer the dominant stimulus (preferred to non-preferred switch; P2NP) are locked to the hyperpolarising phase (−147.6°, starting at ∼750ms before the switch, Figure 6D). This selective locking could result in the modulation of the membrane potential, thereby pushing the two ensembles closer to, or farther away from the firing threshold, thus increasing or decreasing the firing probability, respectively.

### Intrinsic nature of prefrontal state fluctuations

If the competition between low-frequency transients and beta-bursts that regulate access to visual consciousness is intrinsically generated, reflecting waking state fluctuations, traces of this process should also be observed during resting-state: i.e., in the absence of any sensory (i.e. visual) input. Indeed, in resting-state, low-frequency bursts suppressed beta activity (Fig. 7A). Periods of uninterrupted beta activity exhibited a gamma distribution with duration of 1.2 ± 1.44s (median±SD), which is close to the psychophysical distribution of stable perceptual dominance durations (1.54 ± 1.28s) (Fig. 7B). Therefore, prefrontal state fluctuations appear to reflect an intrinsic process observed in the macaque PFC.

**Figure 7.**
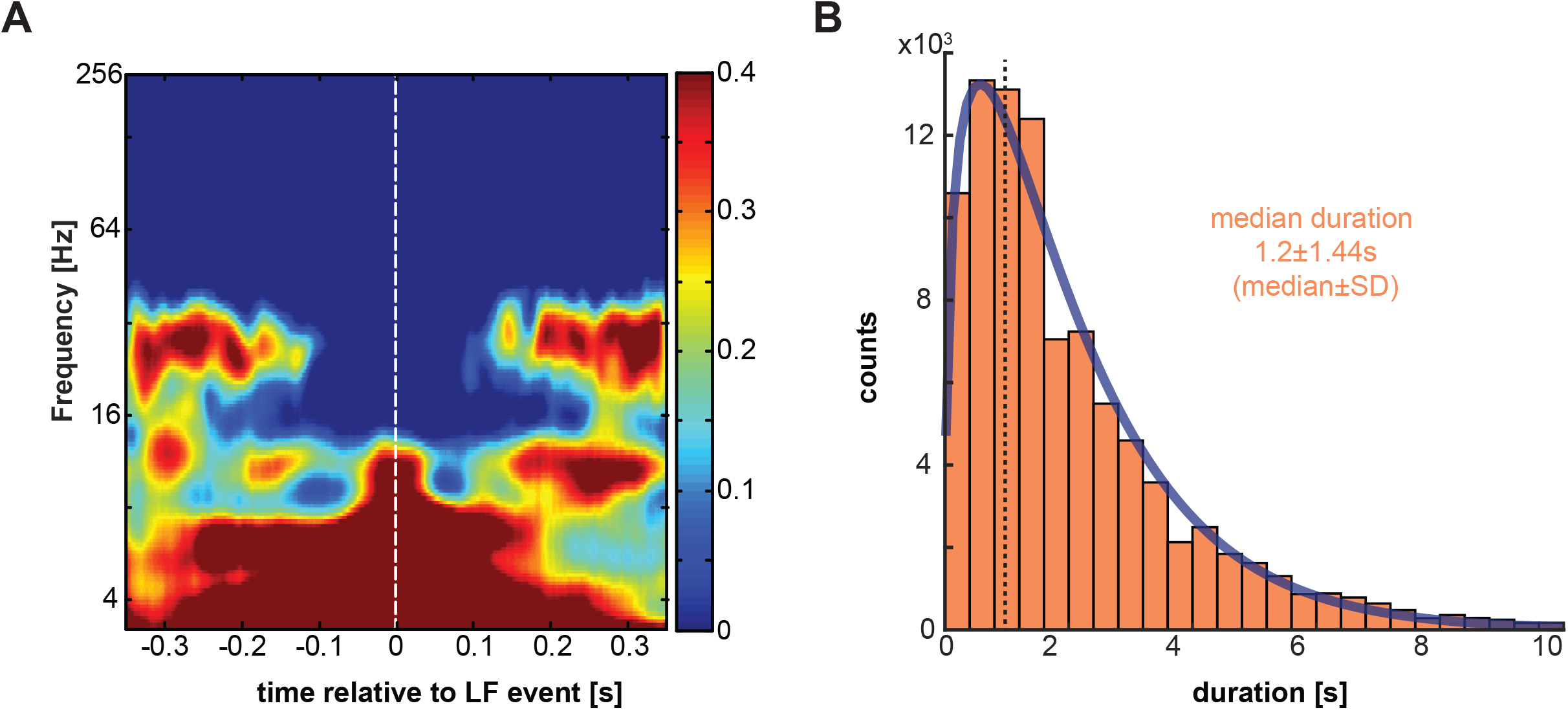
Intrinsic states during resting state show dynamics similar to perceptual transitions. A. Grand-average (n=480), low frequency burst-triggered spectrogram during resting state showing cortical state fluctuations between low frequency and beta activity in the absence of structured sensory input. B. Periods of sustained beta activation (1.2 ± 1.44s, median±SD) follow a gamma distribution (BIC (Bayesian information criterion (Schwarz, 1978)) = 2.3 x e^105^ for a gamma distribution vs BIC = 2.9 x^^105^ for an exponential) during resting state (activity periods longer than 10s were discarded to maintain equivalence with the experimental trial durations). The median duration is remarkably similar to the psychophysical gamma distribution (1.54 ± 1.28s, median±SD).

## Discussion

Contrary to the notion that PFC activity could reflect only the consequences of conscious perception (Boly et al., 2013, 2017; Frässle et al., 2014; Koch et al., 2016; Lamme, 2006; Sandberg et al., 2016; Tsuchiya et al., 2015), our results suggest that perisynaptic prefrontal state fluctuations precede spontaneous changes in the content of consciousness, with the latter reflected in the activity dominance or suppression of feature specific prefrontal ensembles, in the absence of any voluntary report requirements. Therefore, in the PFC there are neural events preceding coding from ensembles, a topic that is significant from a theoretical point of view in order to understand the cortical organisation of conscious processing (Maillé and Lynn, 2020; Seth, 2018).

Previous electrophysiological studies in the PFC have revealed representations of the content of consciousness using exogenous perceptual manipulations and behavioral reports (Panagiotaropoulos et al., 2012; van Vugt et al., 2018); (Gelbard-Sagiv et al., 2018; Libedinsky and Livingstone, 2011; Panagiotaropoulos et al., 2012; van Vugt et al., 2018) as well as preparatory spiking activity before spontaneous perceptual changes (Gelbard-Sagiv et al., 2018; Libedinsky and Livingstone, 2011), that could not however be dissociated from the signals related to voluntary motor reports. Our findings during a no-report BR paradigm could be associated with a gating-like mechanism (Hazy et al., 2007; O’Reilly, 2006; Rougier and O’Reilly, 2002) where intrinsically-generated fluctuations in the prefrontal cortical state between low-frequency transients and periods of sustained beta-bursts, gate the access of competing perceptual representations to consciousness. While low-frequency transients are associated with a perceptual update process, preceding spontaneous changes in the content of consciousness, beta oscillations are prevalent during periods of perceptual content stability in no-report BR.

### Low-frequency activity build-up and relation to accumulation of evidence

We found that a gradual spatiotemporal build-up in low-frequency activity, shown to be generated internally, leads to a suppression of beta activity, before a spontaneous reversal in the content of conscious perception. This build-up of 1-9 Hz bursts before spontaneous perceptual transitions is reminiscent of the Readiness Potential or *Bereitschaftspotential*; i.e., a steady accumulation of activity in the Anterior Cingulate Cortex (ACC) and Supplementary Motor Area (SMA) preceding the awareness of the volition to initiate a report (Brunia et al., 2011; Kornhuber and Deecke, 1965; Libet et al., 1983; Schurger, 2018; Schurger et al., 2012). This motor-related process is reflected in a spiking activity build-up in these cortical areas before the voluntary motor report of a perceptual transition in BR (Gelbard-Sagiv et al., 2018). Such build-up of activity has been suggested to be an “integration device” to accumulate either ongoing spontaneous, or stimulus-driven activity, therefore being crucial to setting the threshold for conscious access (Moutard et al., 2015; Pereira et al., 2021), or contribute to transitions in binocular rivalry, and its subsequent decision (Cao et al., 2021). Our findings suggest that even in the absence of active subjective reports of conscious perception, this build up is readily observed in the low frequency activity of PFC before spontaneous changes in the content of consciousness.

### Beta oscillations reflect stability of conscious perception

Periods of beta-burst suppression are generally thought to reflect a decrease in endogenous cortical processing, since beta oscillations are suppressed during cognitive processes like attention, decision-making and movement-planning (Alayrangues et al., 2019; Engel and Fries, 2010; Ray and Cole, 1985; Tzagarakis et al., 2010). Beta activity could therefore reflect an intrinsic mode of cortical operation that shields ongoing behavioural and processing states (status-quo) from interference and distractors (Alayrangues et al., 2019; Engel and Fries, 2010; Ray and Cole, 1985). As a consequence, transient decreases in cortical beta activity could increase sensory information relay (David et al., 2015; Miller et al., 2018; Panagiotaropoulos et al., 2014; Spitzer and Haegens, 2017) providing a mechanism for controlling bottom-up sensory processing through top-down knowledge (Miller et al., 2018). Indeed, a recent study (Karvat et al., 2020) in rats showed that spontaneous beta-activity reduced the true positive detection rate of a vibro-tactile stimulus. It appears that the stronger the burst-rate of ongoing beta, the less likely it is for the animal to detect a stimulus. Online detection of the burst-rate and the thus-informed adjustment of the strength of the stimulus in this study revealed that the inhibitory “hold” acts as a gate that requires sufficient stimulus-level push to be opened.

Interestingly, desynchronised brain states were recently suggested to mediate low-level awareness in a theoretical study (Zerlaut and Destexhe, 2017). Our findings are in agreement with such a model since we found that dissolution of beta-coherent dominant ensembles from low-frequency transient bursts precedes intrinsically generated changes between the two perceptual states. This result is reminiscent of rule-selective prefrontal ensembles that are coherent in the beta band with low-frequency activity inhibiting a rule that is about to be deselected (Jensen and Bonnefond, 2013). This suggests that the underlying prefrontal mechanism for the emergence of conscious perception and cognitive control might be the same, relying on desynchronisation of neural activity and supporting a cognitive role for consciousness.

### Prefrontal ignition and gating of access to consciousness

We propose that the disruption of intrinsically-generated beta activity by low-frequency transients could reflect the neural correlate of the prefrontal ignition mechanism that has long been hypothesised to gate or control access to consciousness (Dehaene and Naccache, 2001; Del Cul et al., 2009; Lau and Rosenthal, 2011; van Vugt et al., 2018). In our recordings, apart from BR, we observed the same beta synchronisation after a physical, externally-induced stimulus change. This desynchronisation was accompanied by a strong visually evoked potential (VEP) in the same low-frequency range, around 200ms after the stimulus presentation. Whereas van Vugt and co-authors show a sustained increase in dlPFC activity after the stimulus becomes reportable (van Vugt et al., 2018), a high-amplitude transient in spiking activity was observed in a binocular flash suppression task, around 200ms after the change in the percept (Panagiotaropoulos et al., 2012). This VEP is akin to large-amplitude signals used to infer the ignition mechanism proposed by the GNW (Dehaene and Changeux, 2011). Hence, our results show that prefrontal gating could be the consequence of such an ignition event. Importantly, we show here that the activity of feature-selective ensembles that follow the active percept, shows a non-linear increase in the range of 130-220ms after the mean low-frequency burst time. This activity is then sustained, followed by a slow decay (Figs. 6A, B), as predicted by the GNW.

Disruption of beta activity from transient low-frequency depolarisation events during BR could provide temporal windows for a reorganisation in the discharge activity of neuronal ensembles that encode the competing representations. Indeed, our results show that the populations which are selective to the suppressed stimulus are locked to the depolarising phase of the low-frequency states. This locking would increase the probability of spiking, thus increasing their firing rates when these sites reflect the dominant stimulus after the switch. Indeed, beta-suppression occurred earlier than the reversal in the population discharge activity encoding the content of consciousness by feature-selective ensembles (data not shown). Furthermore, the burst-rate in low-frequencies and concurrent beta suppression was very weak when no switches occured or a dominant percept transitioned to a piecemeal. This indicates that sufficient accumulation of low-frequency activity and its spread to a significant number of neuronal sites, thus igniting them, is critical for spontaneous perceptual reversals. In addition, it appears that the driver of perceptual reorganisation and update is not the spiking activity of selective neuronal ensembles; rather it is a global state signal (the neurons only seem to report the active percept). This top-down mechanism of perceptual reorganisation is fundamentally different from bottom-up mechanisms proposing that competition between monocular neurons in the primary visual cortex (V1) resolves BR (Blake, 1989; Leopold and Logothetis, 1996). Neuronal activity in V1 is indeed only weakly-modulated during BR in both monocular and binocular neurons (Leopold and Logothetis, 1996), while BOLD modulation of V1 is detected in superficial layers, suggesting feedback from higher cortical areas (Qian et al., 2018). Finally, optical imaging signals in V1 during BR can also be observed during anaesthesia (Xu et al., 2016), indicating that V1 activity is not alone sufficient for conscious visual perception.

## Conclusion

Spontaneous cortical activity can attain various states during wakefulness, and reflect sensory-driven activity (Mazzucato et al., 2015; McGinley et al., 2015; Tsodyks et al., 1999). We observed that the suppression of ongoing beta bursts by low-frequency transients, observed during BR and preceding changes in conscious perception, was also manifested during periods of resting state. This suggests that the source of spontaneous transitions in the content of consciousness may be traced to the interaction of visual input with ongoing, waking state fluctuations in the PFC. Taken together; our results reveal a potential role of prefrontal state fluctuations in a gating process that mediates the emergence of conscious perception.

## Acknowledgments

We thank Dr. Yusuke Murayama and the other technical and animal care staff for excellent technical assistance, Prof. Dr. Nicho Hatsopoulos for help with the implantations of the Utah arrays, Prof. Dr. Stanislas Dehaene for his inputs and insights, and Mr. Akshat Jain for his assistance in implementing code for the resting state analysis, and in the preparation of publishable quality figures. We would also like to thank Dr. Michel Besserve, Prof. Dr. B S Dwarakanath and Dr. Majid Khalili-Ardali, for helpful discussions and, in editing and proof-reading the manuscript.

## Author contributions

Conceptualisation: AD, VK, TIP (lead), NKL; Data curation: AD (lead), VK and JW; Formal analysis: AD (lead), VK, JW, LAF; Funding acquisition: NKL; Investigation: AD (equal), VK (equal), TIP (supporting); Methodology: AD (equal), VK (equal), JW & SS (supporting), TIP (equal); Project administration: TIP; Resources: JW, NKL (lead); Software: AD (lead), VK, JW, LAF & SS (supporting); Supervision: TIP; Visualisation: AD (lead), TIP (supporting); Writing - original draft: AD, TIP (lead); Writing - review & editing: AD, VK, LAF, SS, TIP (lead), NKL.

## Methods

### Electrophysiological recordings

We performed extracellular electrophysiological recordings in the inferior convexity of the lateral PFC of 2 awake adult, male rhesus macaques (*Macaca mulatta*) using chronically implanted Utah microelectrode arrays (Maynard et al., 1997) (Blackrock Microsystems, Salt Lake City, Utah USA). We implanted the arrays 1 - 2 millimetres anterior to the bank of the arcuate sulcus and below the ventral bank of the principal sulcus, thus covering a large part of the inferior convexity in the ventrolateral PFC, where neurons selective for direction of motion have been previously found (Hussar and Pasternak, 2009; Safavi et al., 2018). The arrays were 4×4mm wide, with a 10 by 10 electrode configuration and inter-electrode distance of 400μm. Electrodes were 1mm long therefore recording from the middle cortical layers. The monkeys were implanted with form-specific titanium head posts on the cranium after modelling the skull based on an anatomical MRI scan acquired in a vertical 7T scanner with a 60cm diameter bore (Biospec 47/40c; Bruker Medical, Ettlingen, Germany). All experiments were approved by the local authorities (Regierungspräsidium, protocol KY6/12 granted to TIP as the principal investigator) and were in full compliance with the guidelines of the European Community (EUVD 86/609/EEC) for the care and use of laboratory animals.

### Data acquisition, spike sorting and local field potentials

Broadband neural signals (0.1–30 kHz) were recorded using Neural Signal Processors (NSPs) (Blackrock Microsystems). Signals from the Utah array were digitised, amplified, and then routed to the NSPs for acquisition. For the offline detection of action potentials, broadband data were filtered between 0.6 and 3 kHz using a second-order Butterworth filter (the filter was chosen such that it allowed a flat response in the passband while contributing the least phase distortion due to its low order, yet having an acceptable attenuation in the stop band, i.e. a roll-off starting at −20dB). The amplitude for spike detection was set to five times the median absolute deviation (MAD) (Quiroga et al., 2004). Spikes were rejected if they occurred within 0.5 ms of each other or if they were larger than 50 times the MAD. All of the collected spikes were aligned to the minimum. Automatic clustering to detect putative single neurons was performed by a Split and Merge Expectation-Maximisation (SMEM) algorithm that fits a mixture of Gaussians to the spike feature data which consisted of the first three principal components (Kadir et al., 2014) (Klustakwik). The clusters were finalised manually using a cut-and-merge software (Hazan et al., 2006) (Klusters). For the analysis of perisynaptic LFP activity, the broadband signal was decimated to 500 Hz sampling rate using a Type I Chebyshev Filter, preserving frequency components up to 200 Hz.

### Visual stimulation and experimental paradigm

Visual stimuli were generated by in-house software written in C/Tcl and used OpenGL implementation. Stimuli were displayed using a dedicated graphics workstation (TDZ 2000; Intergraph Systems, Huntsville, AL, USA) with a resolution of 1,280 × 1,024 and a 60 Hz refresh rate. An industrial PC with one Pentium CPU (Advantech) running the QNX real-time operating system (QNX Software Systems) controlled the timing of stimulus presentation, and the digital pulses to the electrophysiological data acquisition system. Eye movements were captured using an IR camera at 1kHz sampling rate using the software iView (SensoriMotoric Instruments GmBH, Germany). They were monitored online and stored for offline analysis using both the QNX-based acquisition system and the Blackrock data acquisition system. We were able to reliably capture the eye movements of the animals by positioning the IR camera in front of a cold mirror stereoscope.

Initially, the two monkeys (A11 and H07) were trained to fixate on a red square of 0.2° of visual angle about 45cm away from the monitors that could be viewed through the stereoscope. This dot was first presented in one eye (the location of the red fixation square was adjusted to the single eye vergence of each individual monkey) and the eye-position was centred using a self-constructed linear offset amplifier. While the monkey was fixating the dot was removed and immediately presented in the other eye. Over multiple presentations, the offset between the two eyes was averaged to provide a horizontal correction factor to allow the two dots to be perfectly fused within the resolution limitations of the recording device (1/100th of a degree). The monkeys were trained to maintain fixation within a window of 2° of visual angle during initiation. After 300ms of fixation, a moving grating of size 8°, moving in the vertical direction (90° or 270°) at a speed of 12° (monkey H) and 13° (monkey A) per second, with a spatial frequency of 0.5 cycles/degree of visual angle and at 100% contrast was presented for 1000-2000ms, in the first five experimental sessions. In the sixth session, 200 random dots at 100% coherence with a limited lifetime of 150ms were presented. This marked the first monocular stimulus epoch in both conditions, viz. Binocular Rivalry (BR) and Physical Alternation (PA). At the end of 1000-2000ms, the second stimulus with the same properties as above but moving in the opposite direction was presented to the other eye. In the BR trials, this marked the “Flash Suppression’’ phase. These two competing stimuli were allowed to rival against each other for a period of 6000-8000ms. In the PA trials, switches in the percept were mimicked by alternatively removing one stimulus based on the mean dominance time computed from the Gamma Distributions (tailored to each monkey’s performance and statistics) acquired during multiple training sessions, and adjusted to be closer to a mean of 2000ms. Free viewing within the +/-8° window, which included the stimulus, elicited the Optokinetic Nystagmus (OKN) reflex concomitant to the perceived direction of motion which served in lieu of a voluntary report, fulfilling the criterion of a “no-report paradigm”. The monkeys were given a liquid reward (either water or juice) at the end of the trial, if their OKN successfully remained within the specified viewing window during the entire duration of the trial. Every successful trial was followed by a 2000-2500ms inter-trial period.

### Detection of spontaneous transitions

The recorded eye-movement signal in the Y-coordinate was first low-pass filtered using a 3rd order Butterworth Filter below 20 Hz to remove involuntary jitter-induced high-frequency noise. A custom GUI written in MATLAB allowed us to manually identify the end of a dominance period and the beginning of the subsequent one. Manual marking (performed by two authors, AD and VK) was necessitated due to the large variability in the shapes that comprised the OKN complex. These events were based on the change in the slope of the slow-phase of the OKN. Such spontaneous switches were identified by the difference in the end of a dominance and the beginning of the next one; specifically, if this difference was less than 250ms (a fast switch). A “clean” transition was designated if the previous dominance and the subsequent one lasted for at least 500ms without being broken. Analogous to subjective reports, we aligned the LFP and the spiking activity at the beginning of the subsequent dominance period. This was performed in the same way for both BR and PA trials.

### Treatment of the LFP data

Firstly, the decimated LFP signal (0.1-500 Hz) around the OKN transitions was decomposed into a time-frequency representation using a Continuous Wavelet Transform (CWT, MATLAB 2016b) with a Morse wavelet of 7 cycles. This allowed us to resolve 169 frequencies from 0.5 to 256 Hz (500 Hz sampling rate) while preserving the full temporal resolution. The CWT for each channel in each transition (BR, PA, PM and RT) was first z-scored in the frequency domain to visualise the relative changes in power and then pooled across all channels and averaged. To visualise the differences between spontaneous transitions, piecemeals and randomly-triggered periods, the latter two spectrograms were subtracted from the former, respectively.

To understand the evolution of the LFP activity, we first filtered the broadband LFP trace into two constituent bands that were identified to be modulated during the task from the time-frequency analysis, i.e., the low-frequency (1-9 Hz) and the beta band (20-40 Hz). We used a 4th and 8th order Chebyshev Type I filter respectively, with a maximum passband ripple of 0.001dB. To get the instantaneous amplitude in time, we transformed the signal into the Hilbert space and then computed the absolute value. Bursting events were detected at each transition in each channel using a threshold which was 4 times the standard deviation of the noise modelled as a Gaussian distribution. The minimum duration of each event to be detected was set as one full cycle of the highest frequency in that band, i.e. 111ms for the 1-9 Hz band and 25ms for the 20-40 Hz band (Logothetis et al., 2012). The event-rate in time was computed as a quasi-PSTH by turning the detected bursts into a binary spike-train and smoothed with a Gaussian kernel of width 25ms, and then averaged across all channels (events/s/transition). The burst rate was computed as the sum of low-frequency bursts normalised by the number of transitions and channels (events/transition/channel). To compute the build-up in the low-frequency activity, the amplitude at each detected time-point was averaged first across all channels for a given transition, and then averaged across all transitions. A line was then fit to this mean scatter-plot using the CurveFit Toolbox in MATLAB.

### Construction of direction of motion specific neural ensembles

Single neuron selectivity was assessed during perceptual transition periods of binocular rivalry (perceptual switches) and physical alternation (stimulus switches). During binocular rivalry trials, these periods were selected according to the following criteria: 1. Perceptual dominance (judged from the OKN signal) must be maintained for at least 1000 milliseconds post a perceptual switch 2. A preceding perceptual dominance for the competing stimulus must be maintained for 1000 milliseconds, and finally 3. The delay between the end and the beginning of the two dominance phases was not more than 250 milliseconds. For physical alternations, we selected trials, wherein a stimulus was presented for at least 1000 milliseconds before and after a stimulus switch. The spiking activity was triggered at the beginning of a forward dominance (BR) and stimulus change (PA).

Selectivity was assessed by comparing the distributions of the total number of spike counts across trials where the upward drifting grating was perceived, post (0 to 1000 ms) or pre-switch (−1000 to 0 ms), with trials where a downward drifting grating was perceived. We used a Wilcoxon rank sum test and all neurons where p<0.05 were considered as selective. For a given transition, spikes were binned in 50ms bins for each selective neuron, and the resultant spike-count histograms were summed across the neurons that make up each selective ensemble to represent a population vector.

To analyse the crossover times between the two competing populations, we computed the trend in these normalised direction-selective population sum PSTH activity for every transition in a 900ms window around the time of the marked smooth pursuit OKN change [−900 to 900] by smoothing the raw ensemble population vectors for the two competing populations using a LOWESS filter. Next, we detected each intersection between these two given vectors using standard interpolation. Where multiple intersection points were detected, only that point was considered which was followed by divergences for a minimum of 200ms before and after the intersection point, denoting distinct encoding of the currently active percept. Finally, all cross-over times either less than −0.75s or more than 0.75s were discarded as these were considered to come from noisy trials.

### Spike-field Coherence

The spike-field coherence (SFC) was computed between the spiking activity of selective ensembles for each transition, and the global LFP for that particular transition averaged over all electrodes. A rate adjustment and a finite-size correction was applied before computing the SFC via a multi-taper method (Bokil et al., 2010) (Chronux Toolbox). The mean angle of spike-LFP locking was computed using the pairwise phase consistency method implemented by FieldTrip Toolbox (Oostenfeld et al, 2011).

### Treatment of resting-state activity

LFPs from two continuously-recorded resting state sessions on days when no task-recording was performed, were decimated to 500 Hz as mentioned above. In each channel, the beta bursts were detected using the previously-mentioned LFP event-detection algorithm. The mean of the inter-event interval was used as a threshold to decide which collection of events constituted a phase of sustained activity. These epochs were collected across all channels and pooled across the two monkeys. Both a gamma and an exponential distribution were fit to the observations, with the gamma distribution clearly revealing lower AIC and BIC measures, thereby pointing to this distribution being a better fit than the exponential.

### Statistical methods

All statistical comparisons were performed using a Wilcoxon ranksum test (Hodges and Lehmann, 1963) due to the non-gaussian nature of the underlying distribution from which the data originated. Distributions were fit using the MATLAB statistical toolbox using a Maximum-Likelihood-Estimate method. For model comparisons, the allFitDist.m toolbox was used that also generated metrics for appropriate model selection. For non-parametric fitting of distributions with widely different sample numbers, the kernel density estimate method implemented in the MATLAB Statistics Toolbox was used to generate the best-fit function, which was then normalised for visualisation. For computing significant phase angles of spike-LFP coupling, the Rayleigh Test for circular data was used.

---

**SI Figure 1.**
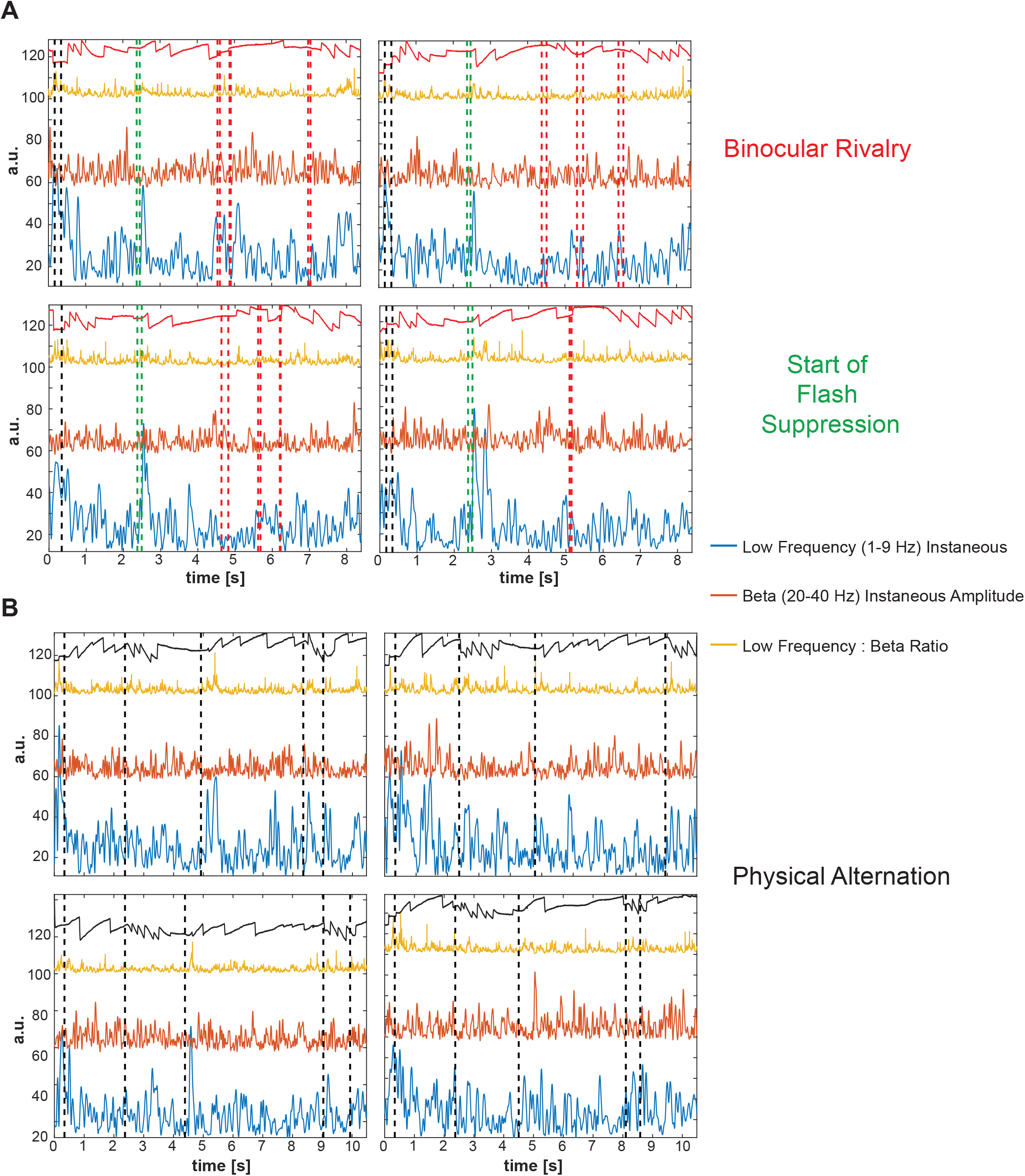
Low Frequency and Beta Instataneous Amplitudes corresponding to a percept switch. A. The low frequency activity averaged across all the 96 channels for a particular transition in Binocular Rivarly(BR) trials shows a large amplitude deflection just before a spontaneous percept switch (marked by the change in the polarity of the Optokinetic Nystagmus(OKN)), along with a concomittant suppression in the beta regime. B. The low frequency activity averaged across all the 96 channels for a particular transition in Binocular Rivarly(BR) trials shows a large amplitude deflection, this time after a physical switch, along with a similar concomittant suppression in the beta regime. This amplitude increase is even greater in magnitude because it is a visually evoked potential.

**SI Figure 2.**
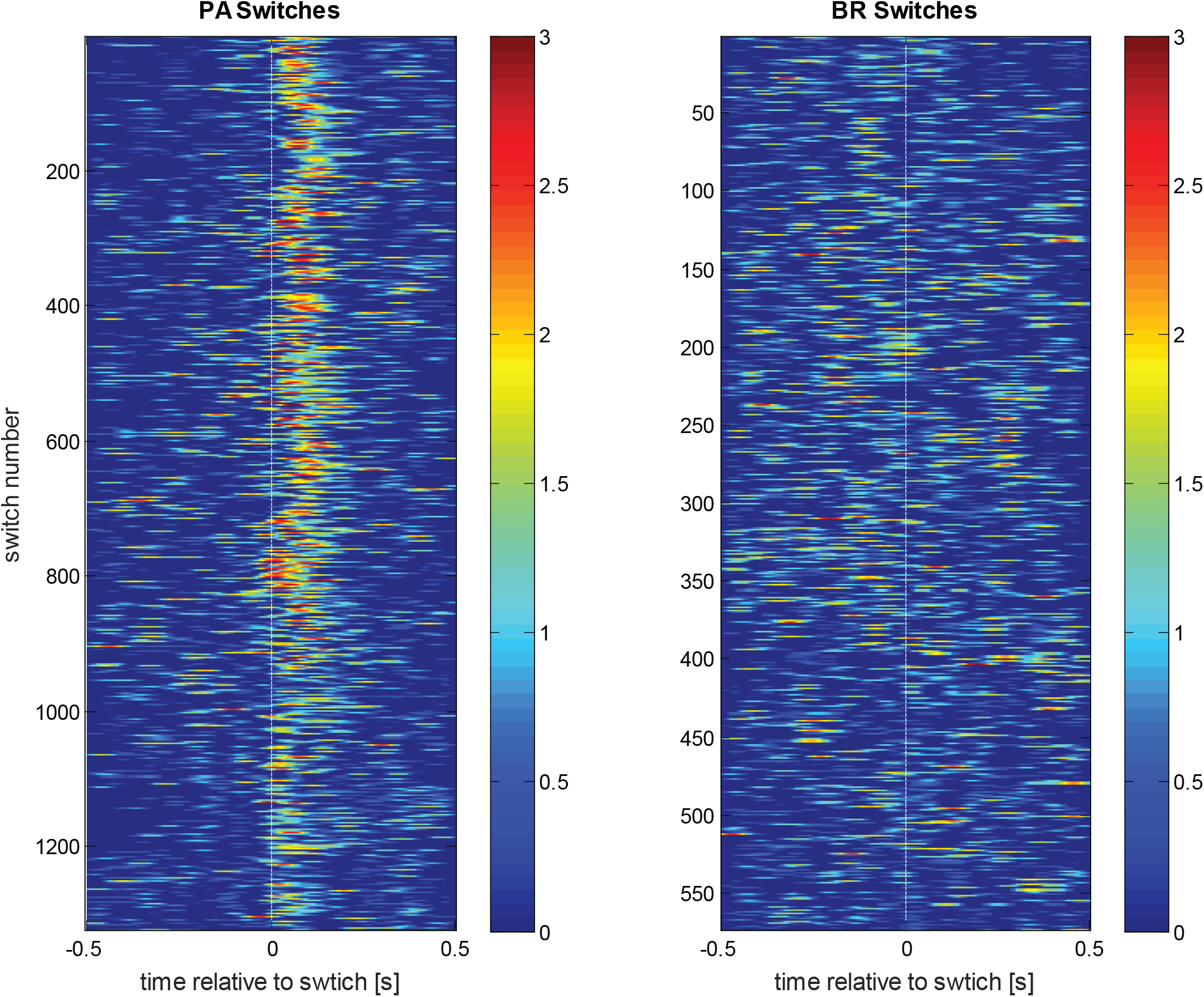
Qualitative analysis of the temporal jitter in the 1-9Hz instantaneous amplitude. This figure depicts the 1-9Hz instantaneous amplitude extracted around clean switches (i.e. a minimum dominance of 500ms before and after a transition) in both the physical alternation (left) and binocular rivalry (right) conditions. While after the transition in PA, a strong and consistent visually-evoked potential is observed, the activity around a spontaneous switch is rather diffuse, yet shows a preponderance of low-frequency activity rising and concentrated in the pre-switch period.

**SI Figure 3.**
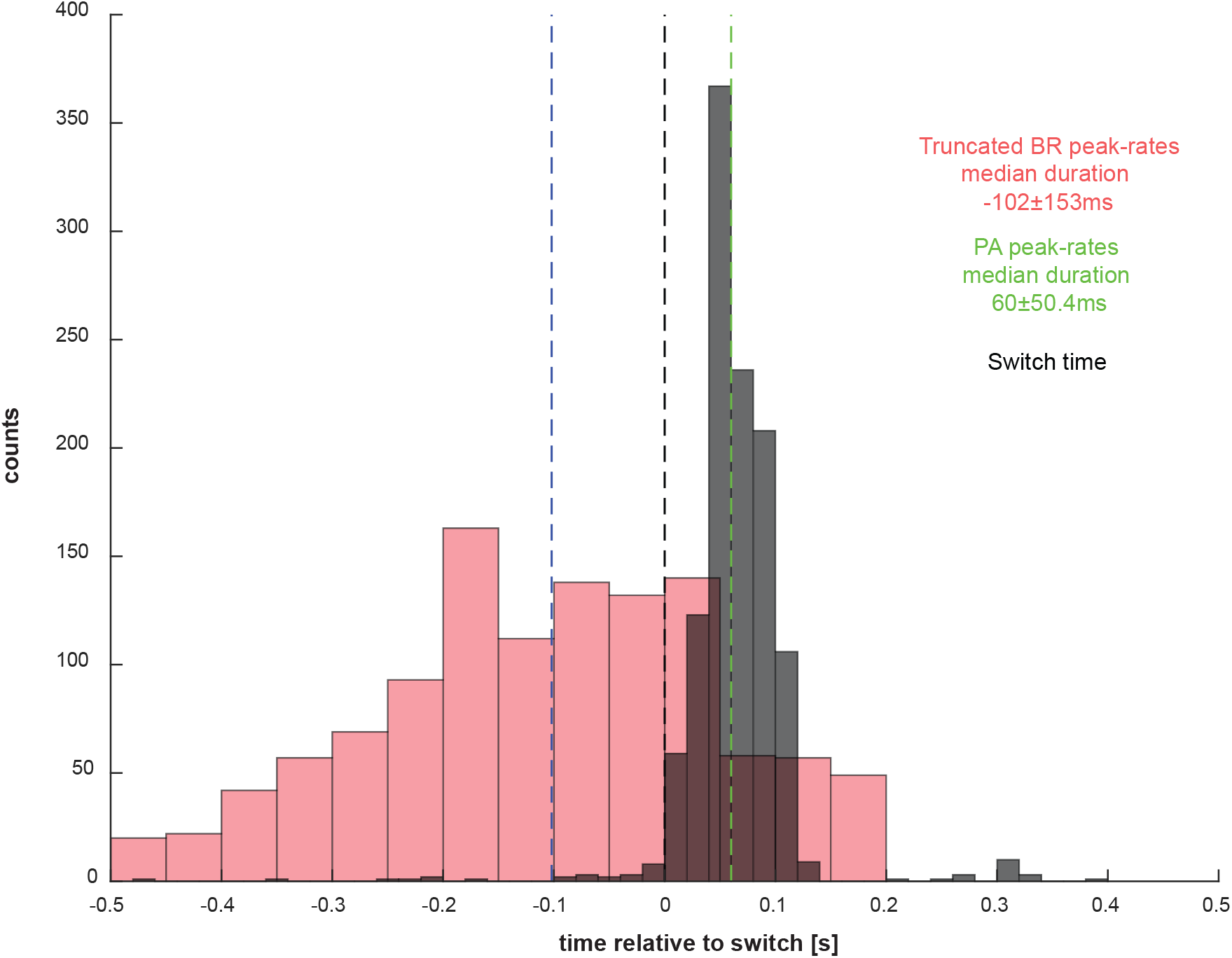
Truncated low-frequency peak rate comparison. Because events towards the end of the post-switch window in BR could signal an upcoming transition, we discarded these events that occurr after 150ms (timing of the end of the VEP in PA). The difference between the median timing of the peak rate in BR and the VEP in PA was further enhanced.

**SI Figure 4.**
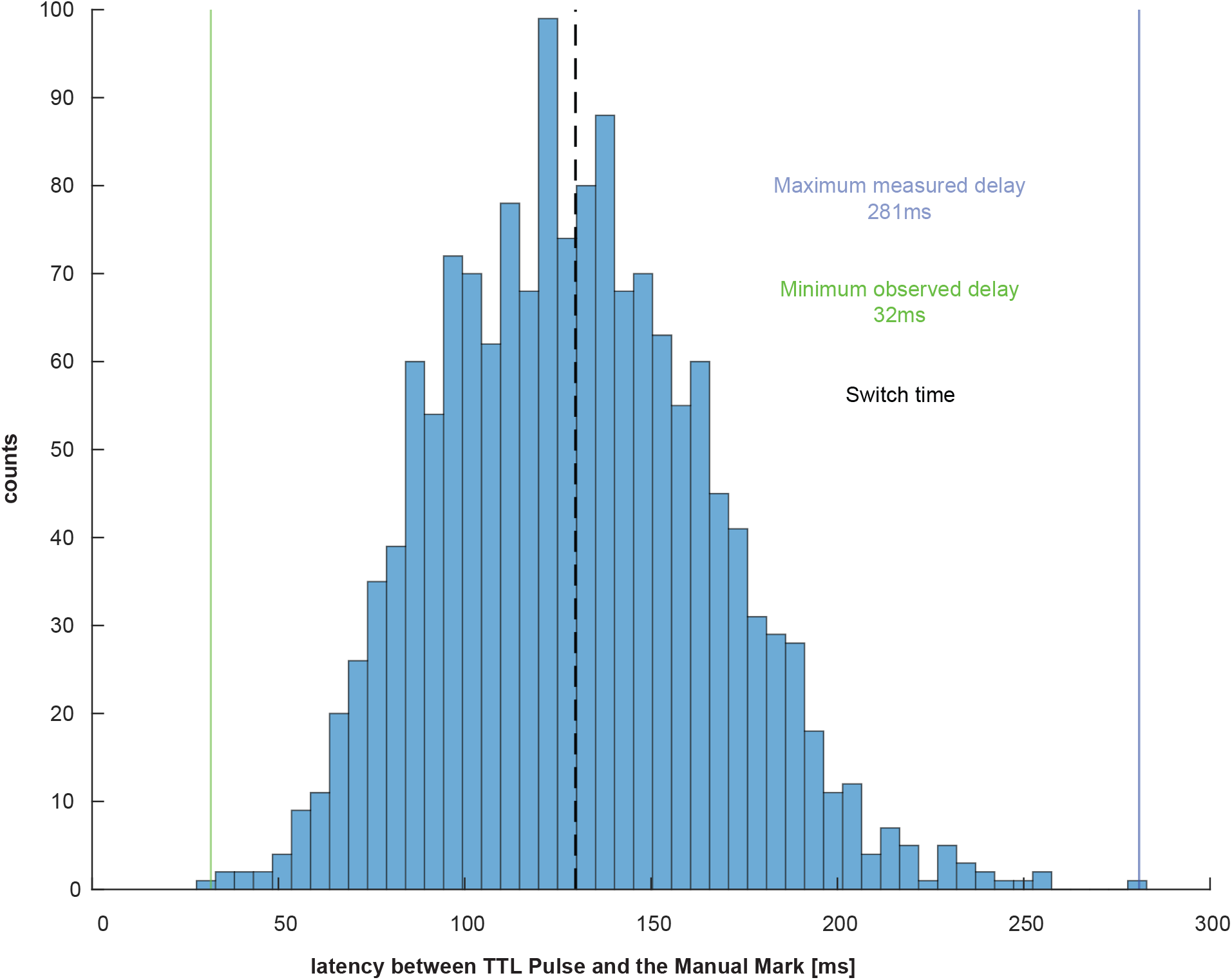
Comparison of the difference between the TTL pulse and the manually marked change in Physical Alternation. We computed the latency between the onset of the subsequent stimulus and the succeeding change in the polarity of the induced OKN for all PA trials. We found an average latency of 129.4±36.5 ms (mean±SD), which indicates that the change in the eye-movement is induced within a very short interval. The minimum latency observed was 32ms whereas the maximum observed latency was 281ms.

**SI Figure 4.**
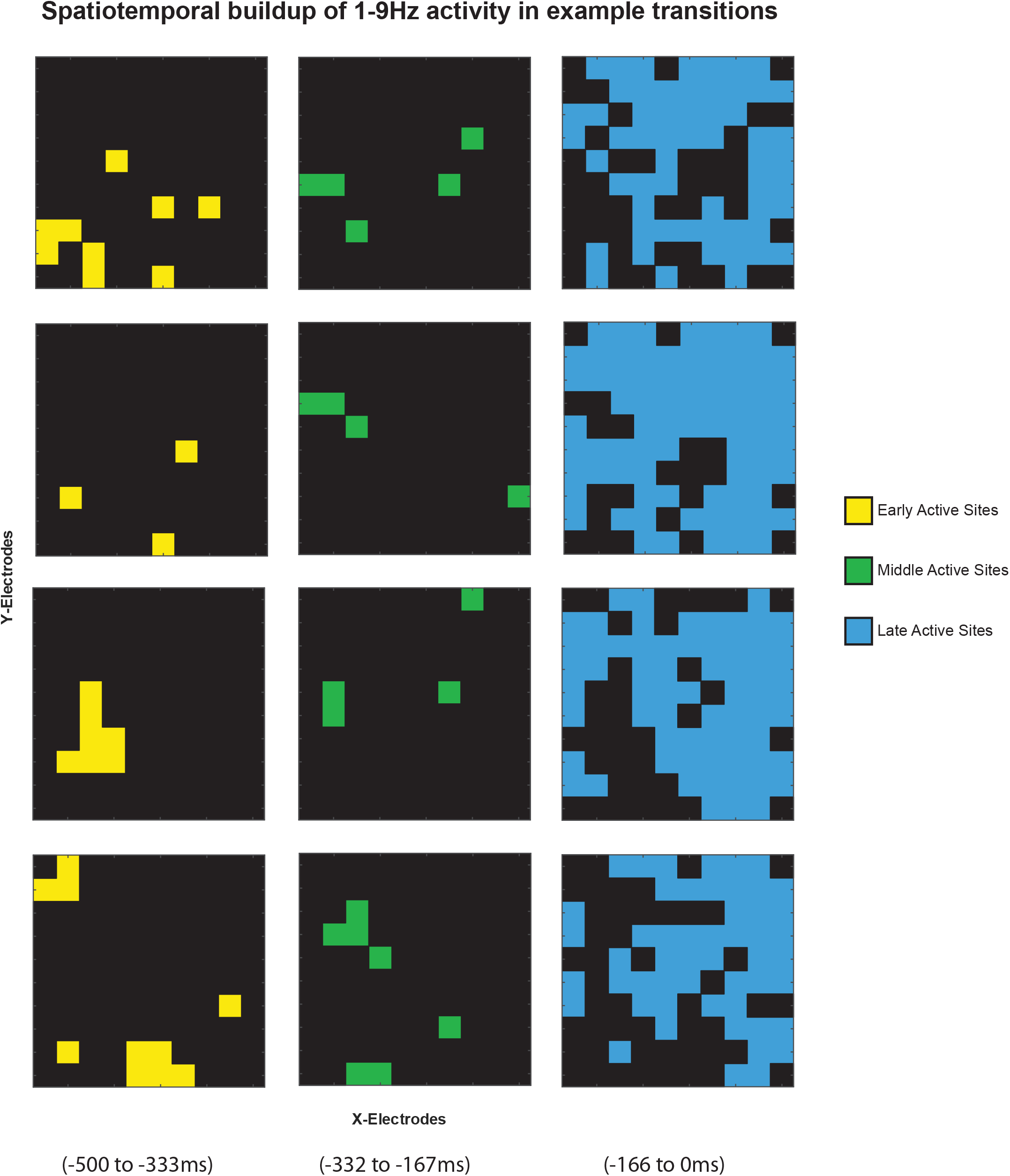
Spatial buildup of low-frequency activity. This figure shows the spatial buildup of 1-9Hz activity in 3 temporal windows viz. in an early window ([−500 to −333ms]), a middle window ([−334 to −166ms]) and a late window ([−167 to 0ms]) preceding four typical spontaneous switches. Progressively, more sites are activated approaching a switch.

**SI Figure 5.**
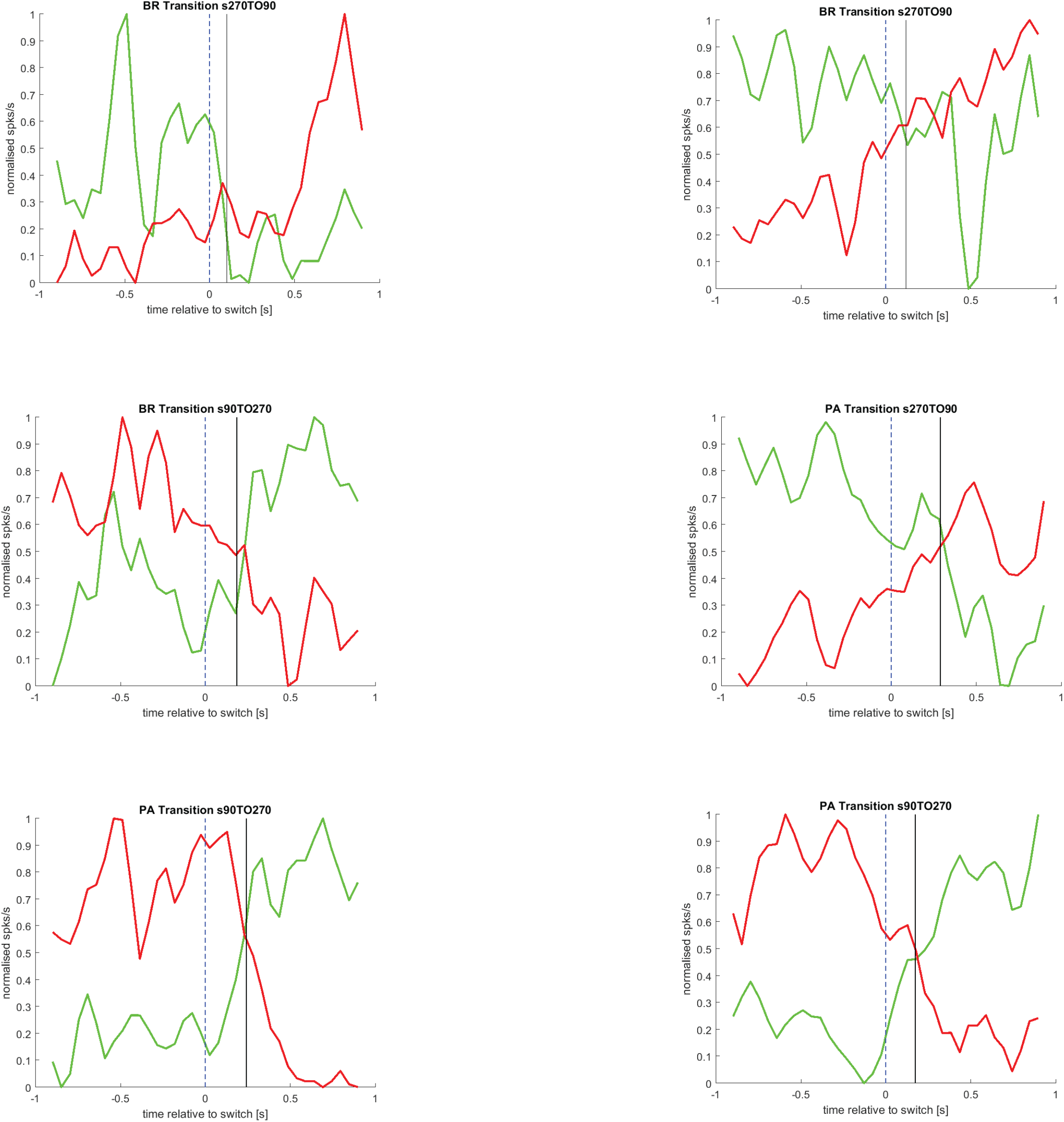
Estimation of change in encoding in selective ensembles. Cross-over points of the normalised population vectors (summed spike counts per bin) describe the point of change in the encoding of the dominant percept by the selective ensembles of neurons. These points are computed by first extracting the best-fit trend of the population vectors using LOWESS smoothing, and then estimating the intersection using interpolation. If multiple intersections are estimated, then that intersection point is chosen such that for 200ms before and after it, there are no other crossings, i.e. stable divergence is observed. Three examples of BR and PA transitions each are shown above (green = downward selective population; red = upward selective population). These cross-over points are then collected across all transitions.

